# Materials in action: The look and feel of soft

**DOI:** 10.1101/2021.01.22.427730

**Authors:** Müge Cavdan, Knut Drewing, Katja Doerschner

## Abstract

The softness of objects can be perceived through several senses. For instance, to judge the softness of our cat’s fur, we do not only look at it, we also run our fingers in idiosyncratic ways through its coat. Recently, we have shown that haptically perceived softness covaries with the compliance, viscosity, granularity, and furriness of materials (Dovencioglu et al.,2020). However, it is unknown whether vision can provide similar information about the various aspects of perceived softness. Here, we investigated this question in an experiment with three conditions: in the haptic condition, blindfolded participants explored materials with their hands, in the visual-static condition participants were presented with close-up photographs of the same materials, and in the visual-dynamic condition participants watched videos of the hand-material interactions that were recorded in the haptic condition. After haptically or visually exploring the materials participants rated them on various attributes. Our results show a high overall perceptual correspondence between the three experimental conditions. With a few exceptions, this correspondence tended to be strongest between haptic and visual-dynamic conditions. These results are discussed with respect to information potentially available through the senses, or through prior experience, when judging the softness of materials.

## Introduction

Objects in our world consists of single or composite materials. To be able to swiftly judge and recognize properties of materials is important, because perceived material qualities influence how we interact with an object. Humans have this ability and are able to make judgments about materials visually and haptically: we move a polished gemstone to visually judge its sparkle and rub a cloth to understand if it is soft enough to wear it. Recent research suggested that perceptions of different aspects of material qualities may be mediated by different senses (e.g., vision or touch, Adams et al., 2016; Sahli et al., 2020). Often, however, also the same aspects of material qualities are judged through different senses: to judge the softness of our cat’s fur, we do not only look at it, but we also run our fingers in idiosyncratic ways through its coat. In this example, the softness of the material can be assessed directly, by touching the cat (Lederman & Klatzky, 1987; Di Luca, 2014; Cavdan et al., 2019; Dovencioglu et al., 2020), - but also indirectly, by looking at it (Bergmann-Tiest & Kappers, 2007; Giesel & Zaidi 2013; Schmidt et al., 2017). What is not known though is, whether these two routes of processing might yield the same evaluations of softness.

Not just softness, but many material qualities are directly available through touch. Indeed, the topic has attracted attentions in haptics community for quite a while (Lederman, 1974; Srinivasan et al., 1990; Srinivasan & LaMotte, 1995; for a review see: Bergman-Tiest, 2010) – increasingly so in the past few years (Cellini et al. 2013; Drewing et al. 2018; Vardar et al. 2019; Mezger & Drewing, 2019; Cavdan et al., 2019; Dovencioglu et al., 2020; Cavdan et al., 2020; see Okamoto et al., 2013 for a review). According to a meta-analysis by Okamoto et al. (2013) the tactual properties of materials can be categorized in five main sections which are warmness (cold/warm), hardness (hard/soft), micro and macro roughness, and friction (moistness/dryness, stickiness/slipperiness).

Visually, only some material properties can be judged directly from images, such as surface gloss, transparency or translucency. Thus, a large majority of research on the visual material perception has centered on those problems (e.g., see Chadwick & Kentridge, 2015 for a review). Softness is related to the subjective impression of the compressibility and deformability characteristics of things and materials, which typically includes a relation to forces that can be directly sensed by touch, but not by vision. However, because of our lifelong experiences with materials, i.e., looking at them while we interact with them, we are also able to judge indirectly material properties from images, e.g., their rigidity, wobbliness or stickiness (Alley et al., 2020; Schmidt et al., 2017). Especially, when we watch objects move and materials deform, impressions of material qualities can be perceived quite vividly (van Assen et al. 2019; Schmid & Doerschner, 2018; Alley et al., 2020; Schmidt et al., 2017; Bi & Xiao, 2016; Doerschner et al., 2011; Mao et al. 2019; Marlow & Anderson, 2016; Yilmaz & Doerschner, 2014; Sakano & Ando, 2010).

Interestingly, it is not just the deformation of a material that triggers a particular impression of the material quality but also watching the interaction with a material: when we actively explore materials in order to gain information about the objects, we adjust our hand and finger motions to the material properties, e.g. we tend to rub rough materials such as felt (Dovencioglu et al., 2020) and to the information we want to gain, e.g. we apply pressure when we wish to find out about an object’s deformability (Lederman & Klatzky, 1987; Lezkan et al., 2018; Cavdan et al., 2019; Zoeller & Drewing, 2020; Cavdan et al., 2020). We know that observers can estimate the weight of lifted objects by just observing the lifting motion (Bingham, 1987; Hamilton et al., 2004; Hamilton et al., 2005; Auvray et al., 2011; Maguinness et al., 2013), and more recent work has shown that humans can distinguish compliance by observing someone else’s finger motions (Cellini et al. 2013; Drewing & Kruse, 2014). Similarly, there has been evidence that visually observing exploratory hand motions of others can yield impressions of material qualities (Yokosaka et al., 2018; Wijntjes et al., 2019).

Do these sources of information, i.e., images, motion/deformations of the material, watching hand movements and haptic exploration^1^, provide rather complimentary or mostly redundant information? A high degree of redundancy might yield quite similar perceptual spaces when estimating material qualities on any of these sources of information (images, haptic, image motion etc.) in isolation. Whereas complimentary processing might yield somewhat different impressions of a material quality, say softness, when elicited by different sources of information. While cue combination studies might provide some important insights into how information is integrated (Lederman et al., 1986; Lacey & Sathian, 2014; Cellini et al., 2013; Adams et al., 2016; Paulun et al., 2019; Wolfe, 1898; Ellis & Lederman, 1999), it is also of interest to understand how much the perception of a material quality from one source of information corresponds to the perception of the same material quality from another source of information. There are only a few studies that have investigated this. For example, Vardar et al. (2019) analyzed similarity ratings for a set of various materials (mounted flat on wood) based on visual or haptic comparisons and found the organization of the perceptual spaces suggests that vision and touch rely on congruent perceptual representations. Baumgartner et al. (2013) used a more extended set of materials, but again limited to textures mounted flat on wood, and assessed ratings of material qualities for visually and haptically presented materials. They conclude that material representations might overall be similar in visual and haptic domains, however how well ratings in visual and haptic domains agree, appears to be depended on the attribute being rated (Baumgartner et al. 2013, their Figure 7). Xiao et al. (2016) investigated the perception of fabrics and found that visuo-haptic matching improves when visually presented fabrics were draped instead of mounted flat, and Wijntjes et al. (2019) showed that movies can reveal more about how fabrics feel than can still images.

What might complicate comparisons of perceptual experience across the senses is that many perceptual attributes are not very well defined. For example, we have recently shown that, perceived *softness,* which, in haptic research, has traditionally been equated with the compliance of a material (Kaim & Drewing 2011; Cellini et al. 2013; Di Luca, 2014; Punpongsanon et al., 2015; Kitada et al, 2019; Zoeller et al., 2019) is in fact a multidimensional construct, that consists of several qualities such as surface softness, granularity, and viscosity (Cavdan et al., 2019; Cavdan et al., 2020; Dovencioglu et al., 2020). This makes intuitively sense: the softness of sand on a beach is different than the softness of a rabbit’s fur, or the softness of an avocado, and we even found that this dimensionality is reflected in the way we explore the material and what property we judge (Cavdan et al., 2019; Cavdan et al., 2020; Dovencioglu et al., 2020). Thus, if one were to compare haptic and visual perception of softness, one would have to be careful to compare all of the underlying dimensions of this perceptual attribute. This is the goal of this study.

Specifically, we seek to understand to what extent the dimensions of perceived softness, that we found in previous haptic experiments, are also present in vision. To do so we conducted an experiment with two visual conditions, including a wide range of materials. In one condition we present movies showing interactions with materials while doing a rating task. This provided observers with the maximum amount of visual information possible, not just showing how materials deform but also typical interactions while rating material qualities (dynamic condition). In a second condition the visual information was reduced, showing only still photographs of the same set of materials (static condition). We compare results of the visual experiment to data from a corresponding haptic study by our group (Cavdan, Doerschner, & Drewing, 2019). We hypothesize that the correlation between the perceptual softness spaces yielded by the two visual conditions should be stronger than the correlation between visual and haptic perceptual spaces, since ratings in the former are based on the same type of indirect information (i.e., visual; Paulun et al., 2017; van Assen et al., 2018; Wijntjes et al., 2020). Given previous results by (Wijntjes et al., 2020), we further hypothesize that the correlation between the perceptual spaces yielded by the dynamic visual condition and the haptic experiment should be stronger than the one between the static visual condition and the haptic experiment.

## General Methods

### Overview

We investigate to what extent the dimensions of perceived softness that we found in previous haptic experiments are also present in visual representation of material qualities. To do this we selected a set of everyday materials that we found to be representative for the various perceptual dimensions of softness in haptic experiments (Dovencioglu et al., 2020; Cavdan et al., 2019). Similarly, we used rating attributes that we found to be strongly associated with the respective perceptual dimensions of softness. Previously recorded hand movements during haptic exploration of these materials were used for the dynamic condition, and still photographs of images of the materials in the static condition. Participants rated all stimuli on all attributes. A Principal Component Analysis (PCA) was used to determine the perceived softness dimensions for static and dynamic visual conditions. We then formally compare the resulting visual and haptic perceptual softness spaces using Procrustes-, and correlation analyses. Along with information pertaining to the visual experiments, we will highlight the relevant methodological and analysis aspects of the previous haptic study (see Cavdan et al., 2019 for more details), which we refer to as haptic condition in the remainder of this paper.

### Participants

Ninety students participated in the experiments: static visual condition: 20 females, 10 males; mean age 23.4 y., range: 20-31y; dynamic visual condition: 21 females; age range: 20-33; mean age:25.1; haptic condition: 21 females, 9 males; mean age: 23.6 years; range: 18-38 years. All of them were right-handed according to self-report, spoke German at a native speaker level and were naïve to the purpose of the experiment. Participants in both visual conditions had normal or corrected-to normal visual acuity and normal color vision (Ishihara, 2004). Participants in the haptic condition had no sensory, motor, or cutaneous impairments and had a two-point discrimination threshold, at the index finger of the right (dominant) hand, of 3 mm or better. Participants provided written informed consent prior to the experiments. All the experiments were approved by the local ethics committee of Giessen University, LEK FB 06, and were conducted in accordance with the declaration of Helsinki (2013).

### Stimuli

Materials items were the same as in our previous haptic study (Cavdan et al., 2019), and included those that had resulted in extreme positive or negative values on four softness-related (Deformability, Fluidity, Hairiness, Granularity) and one control dimensions (Roughness), but included also those that did not show either extreme positive values in any perceptual dimension or that showed extreme values in more than one dimension (see Figure 1, and Dovencioglu et al., 2020, Cavdan et al., 2019).

**Figure 1.**
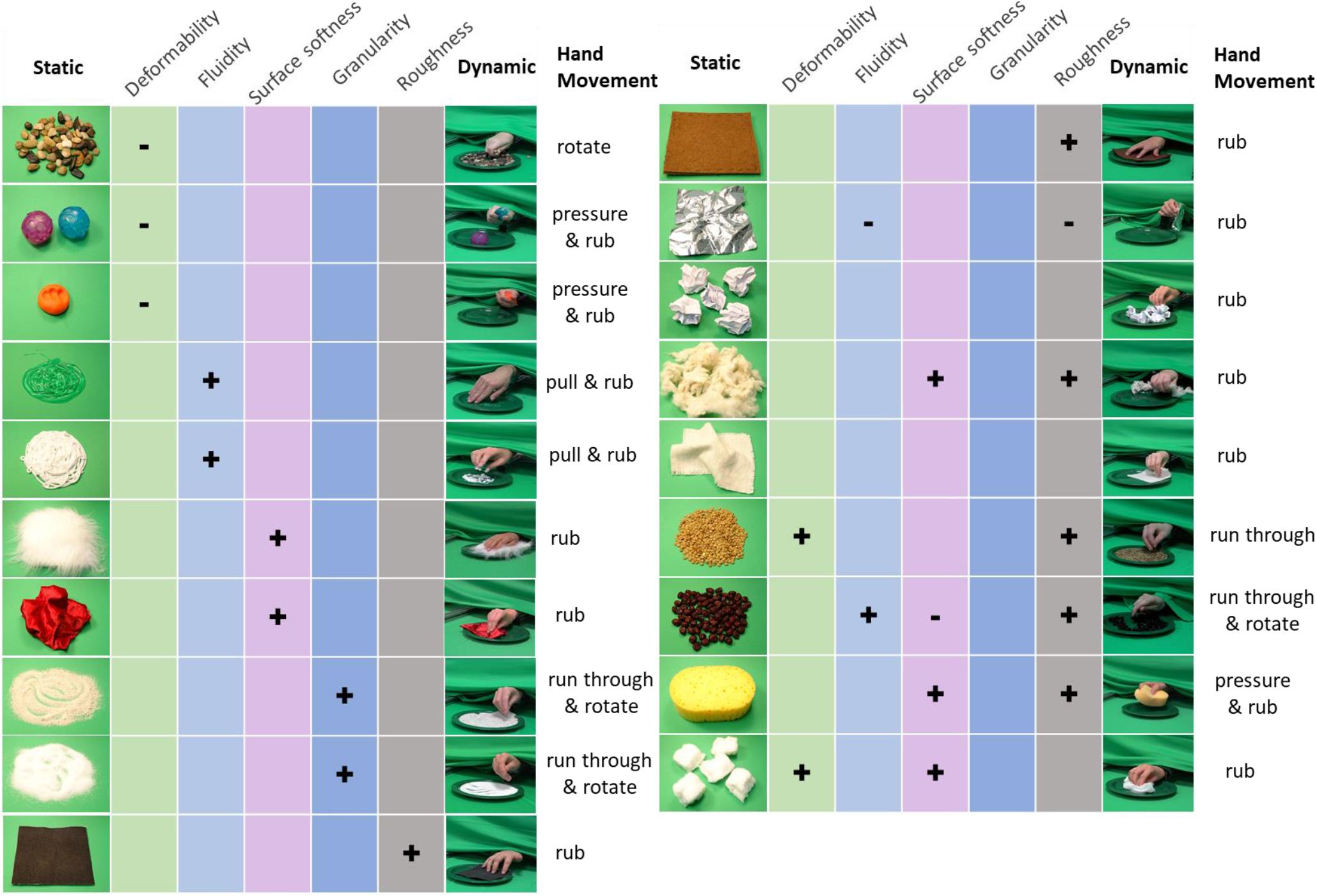
Images used as static stimuli (column 1), associated dimensions of the materials (negative loadings (−), positive loadings (+)), associated hand movements from Cavdan et al. 2019 (column 7), and example frames from dynamic stimuli (column 8). The names of the materials from top to bottom are pebbles, stress balls, play dough, hair gel, hand cream, fur, velvet, sand, salt, sandpaper, felt, aluminum foil, paper balls, wool, linen, lentils, cranberries, sponge, and cotton balls.

#### Still images

To generate still photographs of all 19 materials we placed individual materials on a green cloth (Figure 1). Where possible we added traces of a manual manipulation (e.g., playdough with indentation of fingers, sand with some run-through marks) in order to increase the available shape cues to the respective material properties. Photographs were taken close-up a using a Sony Digital 4K Video Camera Recorder which took 60-bit images at a spatial resolution of 3840 × 2160 pixels (white balance disabled), and with materials illuminated by two 1320 lumen light bulbs placed left and right to the material. The white balance was turned off. This setup yielded a quite natural look, minimizing any harsh shadows. Postprocessing of images centering of the material and cropping to a size of 2049 x 1464 pixels (The Gimp development team, 2019).

#### Dynamic stimuli

For the dynamic condition, we used some of the previously recorded hand movements in the haptic experiments. We selected movies as follows:

First, we determined the either one or two most frequently used, typical exploratory hand movements per material using the taxonomy of by Cavdan et al. (2019, 2020). For example, the most frequently used hand movements for salt were “*run through*” and “*rotate*” (*run through*: “Picking up some parts/portion of the material and letting them trickle through the fingers.”, *rotate*: “Lifting parts of the material to move and turn its boundaries typically inside the finger(tip)s.”; Cavdan, et al., 2019, page 2). Definitions of all exploratory hand movements can be found in Supplementary Methods 1. Most frequently associated hand movements for each material can be found in Figure 1, column 8.

Second, from the movie material collected during the haptic experiment, we selected videos of different participants that performed these typical hand movements. For each of the 19 materials we randomly choose videos of 3 different participants performing the same hand motion, in order to avoid perceptual biases due to a given participant’s potentially unique exploration style. Videos were clipped to 6 seconds (180 frames) with a resolution of 1012 × 1080 pixels. This resulted in 3 matched sets of 19 clips each (one clip per material). Figure 1, column7 shows sample snapshots from the movies used in the dynamic condition.

#### Adjectives

Stimuli were rated on the same 15 sensory adjectives that we used in the previous haptic experiment (Cavdan et al., 2019). These adjectives had been selected based on their association (positive or negative) with the above described softness dimensions or the control dimension (Dövencioglu et al., 2020). These were *soft, elastic, hard, inflexible, moist, wobbly, sticky*, *sandy, powdery, granular, velvety, fluffy, hairy, rough* and *smooth* (see Cavdan et al., 2019 for more details of the selection criteria for adjectives).

### Apparatus

In the static condition, stimuli were displayed on a Samsung UHD (U32D970Q) 32” Professional LED monitor (resolution: 3840 × 2160, refresh rate: 55 Hz). Participants were seated at a distance of about 70 cm from the screen, thus the stimulus size in visual angle on the screen was about 24° in width and about 20° in height.

In the dynamic information condition, stimuli were presented on a DELL UltraSharp monitor (resolution: 2560 × 1440, refresh rate: 56 Hz). Participants were seated at a distance of about 70 cm from the screen, thus the stimulus size in visual angle on the screen was about 15° in width and about 15° in height. Videos were played at a rate of 30 frames per second.

The experiment was programmed in MATLAB 2017a (MathWorks Inc., 2007) using Psychtoolbox routines (Kleiner, Brainard, & Pelli, 2007; Brainard, 1997). Responses were collected with a keypad.

In the previous haptic experiment (Cavdan et al., 2019), we used a curtain to hide the materials from the participant’s view and active noise cancelling headphones so eliminate any contact sounds. Material stimuli were presented on a plastic plate and the participant’s arm was placed on a wrist rest that allowed to explore the materials comfortably from a defined position. Hand movements of the participants were recorded and used in the dynamic visual condition as described in the sections above (for more details of this study please see Cavdan et al., 2019).

### Design and Procedure

#### Static condition

On each trial, participants first saw the to-be-rated adjective. After pressing the space button an image of a material appeared and stayed for 2 seconds at the center of the screen. After the image disappeared observers gave their ratings using the keypad. The task was to indicate how much a given adjective applies to the just seen material on a 5-point Likert-scale item ranging from 1 (adjective *not applicable*) to 5 (adjective *strongly applies*). Participants completed 285 trials (19 materials x 15 adjectives) in about one hour. The order of material-adjective pairs was randomized, every participant saw every material-adjective pair only once (Figure 2).

**Figure 2.**
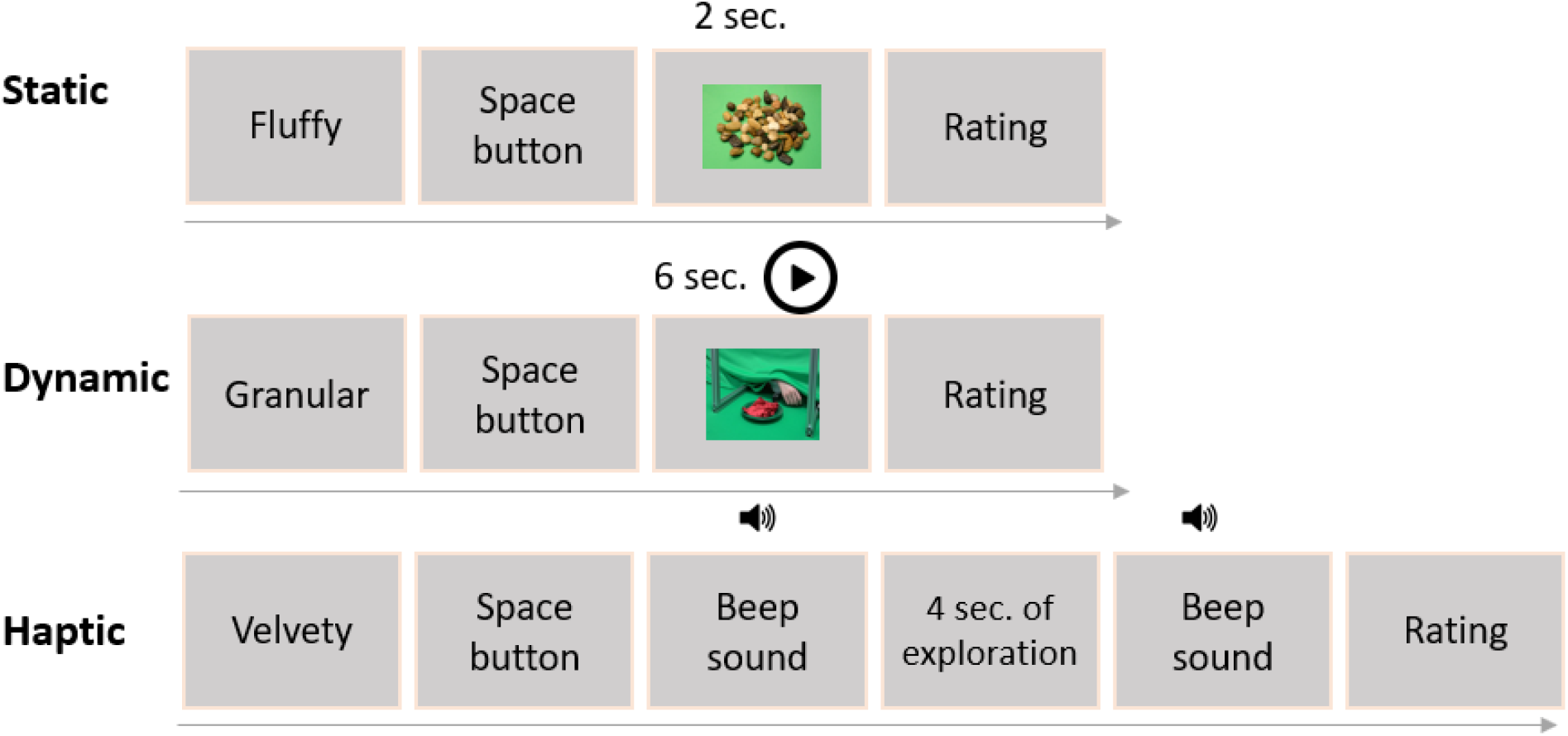
Time course of a trial across conditions in the experiment. First, in all conditions, the adjective to be rated appeared on the screen. After pressing the space button an image was presented on the screen for 2 seconds in static and a video presented on the screen for 6 seconds in the dynamic condition. In the haptic condition a sound (beep) signaled the start for 4 seconds of exploration, and a second beep signaled the end of the trial. Subsequently, in all conditions, participants indicated how much the given adjective applied to the material on a 5-point Likert scale.

#### Dynamic condition

The procedure in the dynamic condition was similar to the static condition except that, instead of a static image, observers now saw a 6 second movie clip showing the exploration of a material (see Figure 2). Observers had to rate all three sets of dynamic stimuli, each set on a separate day. Each session (19 movie clips x 15 adjectives) took about 1 hour.

#### Haptic condition

A typical trial in the haptic condition is shown in Figure 2. In summary, participants first saw the to-be-rated adjective, then pressed the space button to start exploration. Materials were explored for 4 seconds with the right hand. After the exploration, participants removed their hands from the material and rated the material according to the adjective by using keypad using with their left hand. The order of materials and adjectives was randomized, and the experiment took about 1.5 hours.

## Analysis

The goal of this study was to determine the dimensionality of visually perceived softness and to compare it to the dimensionality of the haptic perceptual space.

As a first step we assessed interobserver consistency in the ratings, and checked whether this was approximately in the same range as that obtained for haptic data. Since we acquired 3 data points for each material-adjective combination in the dynamic condition (i.e., three videos for each material-exploration stimulus) we used the average of these 3 scores in the consistency- and all subsequent analyses. Next, we performed separate PCAs for static and dynamic conditions based on average observer data from material-adjective pairs (19 materials × 15 adjectives = 285 data points). Comparing the resulting factor structure and loadings would allow for a first assessment of similarities between visual softness dimensions, and for comparing these to the previously determined haptic perceptual space. To formally assess the degree of similarity conducted a Procrustes analysis on the Bartlett score values of each material across conditions. Should the visual perceptual spaces turn out to be overall similar to each other and the haptic space, we would follow this analysis up with a combined PCA on ratings of the visual conditions and the previous haptic experiment. This would allow for a more fine-grained assessment of the structural similarities of static, dynamic and haptic perceptual softness spaces, e.g., by inspecting the correlations of the respective Bartlett scores between spaces, for all softness dimensions (e.g., static dimension 1 vs haptic dimension 1: static dimension 1 vs haptic dimension 2 etc.).

Lastly, we directly investigated whether there were significant rating differences between the two visual conditions and the haptic condition. To this end, we first calculated mean ratings across participants for each material - adjective pair for the two visual conditions and the haptic experiment, and then computed the distances between these means (3 distances: static-dynamic, static-haptic, dynamic-haptic, for each of the 285 material-adjective pairs). Then we resampled these data using Monte Carlo methods (Efron, 1979), creating a random sample of 10.000 rating distances. Finally, we determined the 95% percentile of this distribution, and report the conditions in which rating difference exceeded this cut-off value.

## Results

### Interobserver consistency

Overall, all interindividual correlations between participants’ ratings in static and dynamic conditions where significant (*p* <.01), and ranged between .41 - .81 and .41-.95, respectively (also see Supplementary Figure 1 & 2). These values were comparable to previously reported results in the haptic condition (.45 - .86) and suggest that participants interpretation of the perceptual meaning of adjectives tended to agree. All subsequent analyses were done on ratings that were averaged across participants.

### PCA for static and dynamic rating data

Since participants showed high consistency in their rating data, we performed next covariance PCAs for static and dynamic conditions. The Keyser-Meyer-Olkin (KMO) criterion was .4, and .5 for the static and dynamic conditions, respectively, which are borderline values. However, Bartlett’s test of sphericity was significant for both conditions (*p*<.01): χ^2^ (105) = 370.32, χ^2^ (105) = 360.03, suggesting that the observed correlations between adjectives were meaningful. Components were extracted according to the Kaiser-criterion and rotated using the varimax method.

In the static condition we extracted three components explaining 83.9% of the total variance (see Table 1). The first component, which we termed *surface softness/deformability,* accounted for 38.2% of the variance with significant loadings of the adjectives *soft, elastic, hard, velvety, hairy, fluffy,* and *inflexible;* the second component, which we named *granularity*, accounted for 25.8% of the variance with significant loadings of the adjectives *granular, sandy, powdery, rough,* and *smooth*; the third component, termed *viscosity*, accounted for 19.9% of the variance with significant loadings of the adjectives *wobbly, sticky, moist.*

**Table 1.** Adjective loadings after rotation for static and dynamic data, as well as for the previous haptic experiment. Each factor labeled based on the adjectives load high (>40% of mean variance per adjective explained: .68 static, .62 dynamic, and .74 for haptic sequentially) and load higher on specific factor than the others. Bold if loading both maximal for adjective and >40% of mean variance per adjective explained, italic if loading only maximal for adjective. Darker colors show positive loadings and lighter colors indicate negative loadings.

| Adjective | Static |  |  | Dynamic |  |  |  | Haptic |  |  |  |  |
| --- | --- | --- | --- | --- | --- | --- | --- | --- | --- | --- | --- | --- |
|  | I. Surface<br>softness/Deformability | II. Granularity | III. Viscosity | I. Surface<br>softness | II. Granularity | III. Viscosity | IV. Deformability | I. Surface<br>softness | II. Granularity | III. Viscosity | IV. Deformability | V. Roughness |
| Fluffy | **1.22** | -0.12 | -0.43 | **1.10** | -0.14 | -0.26 | -0.25 | **1.34** | -0.28 | -0.41 | -0.28 | 0.16 |
| Soft | **1.17** | -0.21 | 0.24 | **0.78** | -0.08 | 0.38 | -0.58 | **0.90** | -0.09 | 0.37 | -0.69 | -0.12 |
| Hairy | **0.84** | -0.10 | -0.41 | **0.83** | -0.10 | -0.27 | -0.15 | **1.07** | -0.39 | -0.25 | 0.06 | 0.48 |
| Velvety | **0.74** | -0.09 | -0.24 | **0.69** | 0.04 | -0.05 | -0.06 | **0.81** | 0.01 | -0.18 | -0.32 | -0.15 |
| Elastic | **0.62** | -0.33 | 0.39 | 0.11 | -0.19 | 0.17 | **-0.75** | 0.10 | -0.32 | 0.28 | **0.80** | 0.09 |
| Hard | **-0.91** | 0.15 | -0.38 | -0.52 | 0.05 | -0.42 | **0.63** | -0.61 | 0.17 | -0.41 | **-0.91** | 0.09 |
| Inflexible | **-0.72** | 0.33 | -0.21 | -0.30 | 0.27 | 0.00 | **0.75** | -0.24 | 0.43 | 0.07 | **-0.84** | -0.04 |
| Sandy | -0.31 | **1.07** | -0.20 | -0.14 | **1.01** | -0.10 | 0.27 | -0.15 | **1.14** | -0.12 | 0.28 | 0.25 |
| Granular | -0.55 | **1.03** | -0.20 | -0.29 | **0.90** | -0.18 | 0.55 | -0.41 | **1.10** | -0.03 | 0.68 | 0.22 |
| Powdery | -0.20 | **0.94** | -0.07 | -0.05 | **0.92** | -0.02 | 0.28 | -0.06 | **0.97** | -0.05 | 0.17 | 0.10 |
| Rough | -0.41 | **0.76** | -0.39 | -0.30 | 0.54 | **-0.57** | -0.02 | -0.39 | 0.38 | -0.33 | 0.14 | **0.70** |
| Smooth | -0.38 | **-0.61** | 0.09 | -0.13 | **-0.50** | 0.23 | 0.30 | -0.27 | -0.20 | 0.11 | 0.12 | **-0.95** |
| Sticky | -0.12 | -0.13 | **.98** | -0.18 | -0.03 | **0.88** | -0.12 | -0.26 | 0.04 | **1.11** | -0.16 | -0.08 |
| Moist | -0.11 | -0.16 | **0.91** | -0.15 | -0.19 | **0.99** | -0.02 | -0.12 | 0.03 | **1.14** | 0.05 | -0.25 |
| Wobbly | 0.18 | -0.27 | **0.78** | -0.13 | -0.16 | **0.70** | -0.42 | 0.03 | -0.22 | **0.96** | -0.40 | -0.04 |

In the dynamic condition we extracted four components explaining 89.2%. While three of the components were rather similar in their structure to the static condition *(surface softness* (25.2%, *soft, velvety, hairy,* and *fluffy*), granularity (23.7%, *granular, sandy, powdery, smooth*), viscosity (21.8%, *wobbly, sticky, moist, rough*)), a fourth component appeared to be exclusively related to the *deformability* of the material. This fourth component accounted for 18.5% of the variance with significant loadings of the adjectives *elastic, hard,* and *inflexible.*

In comparison, in our previously reported haptic condition, we had extracted four components related to softness (*surface softness* (25.9% *soft, velvety, hairy,* and *fluffy), viscosity (*20.6%, *wobbly, sticky, moist*), *granularity* (20.6%, *granular, sandy, powdery*), *deformability* (17.8%, *elastic, hard, inflexible*)) as well as one component related to the roughness of the material (*roughnes*s, 9.5%, *rough, smooth*).

While there are some differences in the number of extracted components between the two visual conditions and the haptic one, it becomes also apparent that there are some structural similarities in the extracted components between the visual and haptic conditions. In particular, inspecting Table 1, in all three conditions, the components of surface softness, granularity and viscosity account for most of the variance in the ratings, which nearly the same patterns of adjective loadings. To formally assess the degree of similarity we conducted a Procrustes analysis on the Bartlett score values of each material across these three components (surface softness, granularity, viscosity) between static, dynamic and haptic conditions. This analysis aims to map two multi-dimensional representations onto each other using linear transformations (reflection, translation, and orthogonal rotation). From this analysis we obtained an index of the error (mean squared error across point pairs) that remains after applying this transform, with lower values indicating better fits. Comparing the mapping of perceptual spaces between the three conditions we obtained values of .19 (static vs dynamic), .20 (static vs haptic), and .25 (dynamic vs haptic). These values were all comparably low (also see Supplementary Figure 4), indicating a rather high similarity between the three perceptual spaces, spanned by the dimensions of surface softness, granularity, viscosity (also see Supplementary Figure 3). Thus, we determined that structural similarity was sufficient to proceed with a combined PCA for static, dynamic and haptic rating data, which would allow us to make more fine-grained comparisons between the static, dynamic and haptic spaces of perceived softness.

### Combined PCA for static, dynamic and haptic rating data

A Keyser-Meyer-Olkin (KMO) value of .68 and a statistically significant Bartlett’s test of sphericity (*χ2*(105) = 1225.62, *p* < .01) suggest that a PCA was indeed appropriate for the combined dataset (Francis & Field, 2011). The combined PCA yielded three components explaining 82.16 % of the total variance. The adjectives *soft, fluffy, hard, velvety, hairy*, *inflexible* and *elastic* loaded high on the first component, which explained 34.5 % of the total variance, and given its loading patterns we named it *surface softness/deformability.* The adjectives *sandy, granular, powdery*, and *rough* loaded high on the second component, which explained 25.5 % of the total variance. We named this second component *granularity.* The adjectives *sticky, moist, and wobbly* loaded high on the third component, which appeared to be related to the viscosity of materials, accounting for 22.16% of the variance. Table 2 shows the adjective loadings for each of the three components in this combined PCA analysis and Figure 3 the corresponding Bartlett scores of all materials, sorted according to their sign and magnitude in the haptic condition, i.e. in order to allow for a better comparison across the three conditions, we kept the ordering of materials in the two visual conditions the same as in the haptic one.

**Table 2.** Rotated Adjective loadings obtained from the combined PCA analysis (static, dynamic and haptic conditions). Color-codes, font styles, and criterion for significance (>40% of mean variance= .67) are as in Table 1.

| Adjective (English /German) | I. Surface<br>softness/Deformability | II. Granularity | III. Viscosity |
| --- | --- | --- | --- |
| Explained variance | 34.54% | 25.46% | 22.16% |
| Soft / weich | **1.06** | -0.20 | 0.37 |
| Fluffy / flauschig | **1.19** | -0.16 | -0.43 |
| Hairy / haarig | **0.86** | -0.12 | -0.43 |
| Velvety / samtig | **0.73** | -0.06 | -0.16 |
| Hard / hart | **-0.85** | 0.22 | -0.48 |
| Inflexible / unbiegsam | **-0.64** | 0.43 | -0.15 |
| Elastic / elastisch | **0.52** | -0.34 | 0.39 |
| Sandy / sandig | -0.23 | **1.08** | -0.12 |
| Powdery / pulverig | -0.14 | **0.94** | -0.03 |
| Granular / körnig | -0.53 | **1.06** | -0.16 |
| Rough / rau | -0.29 | **0.63** | -0.40 |
| Smooth / glatt | -0.37 | **-0.50** | 0.17 |
| Sticky / klebrig | -0.13 | -0.07 | **0.98** |
| Moist / feucht | -0.15 | -0.14 | **0.96** |
| Wobbly / wackelig | 0.14 | -0.25 | **0.85** |

**Figure 3.**
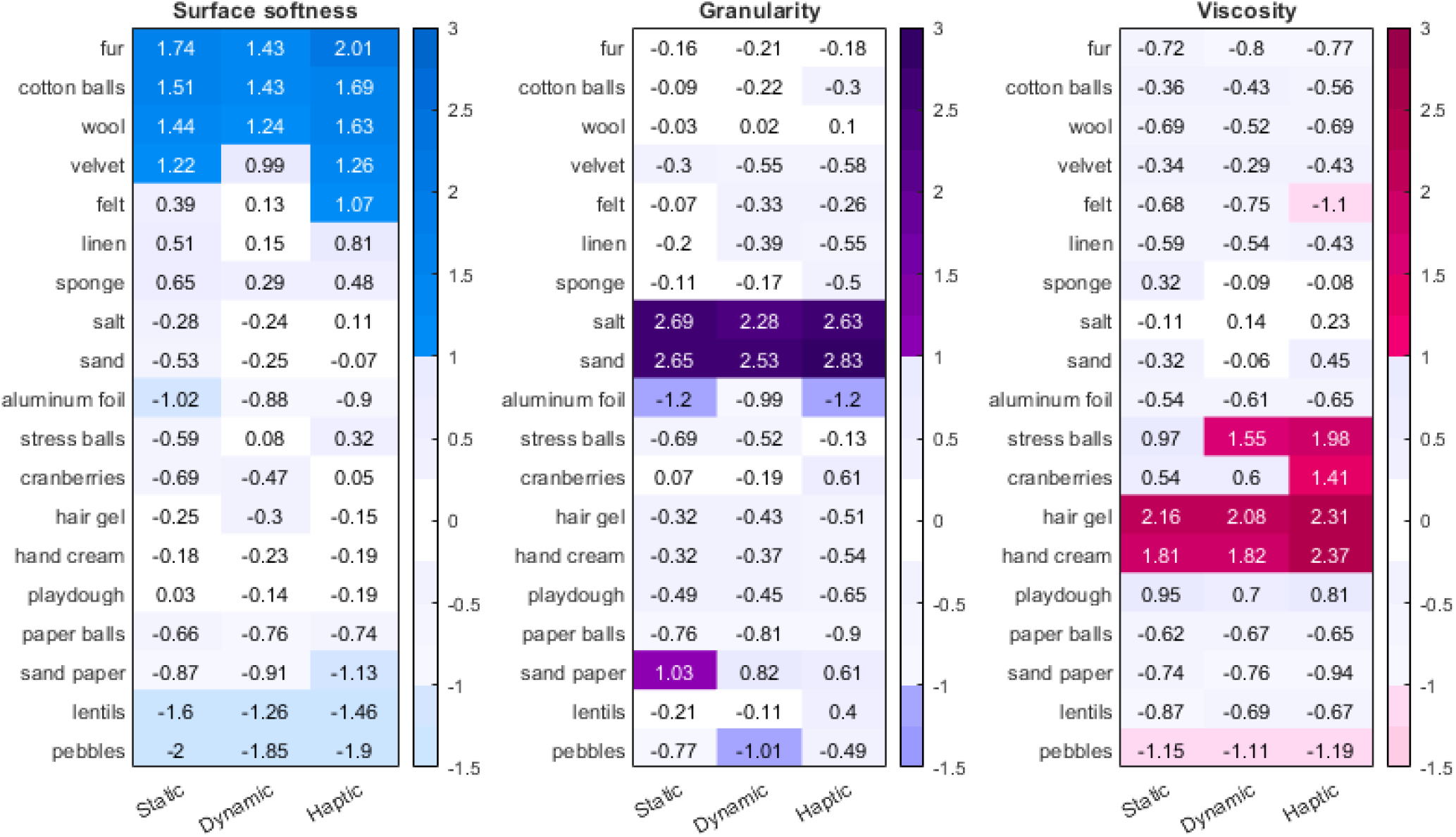
Rotated component scores (Bartlett scores) of materials - in each perceptual softness dimension: surface softness, granularity, and viscosity dimensions - for haptic, dynamic, and static conditions, respectively. Darker, saturated colors indicate positive loadings and desaturated, lighter colors represent negative loadings. Light violet and white areas indicate that loadings were larger than –1 standard deviation, or smaller than 1 standard deviation.

### Assessing similarities between the static, dynamic and haptic spaces of perceived softness

In order to determine the similarities between the static, dynamic, and haptic perceptual spaces of perceived softness, we compared the correlation scores of the Bartlett scores of the three softness dimensions (*surface softness, granularity, viscosity*) across all materials (Figure 3). Figure 4 shows that overall, these correlations were high between the three perceptual spaces, with the highest correlation between the two visual spaces (static & dynamic, (r^*2*^=.95 *p* <.01), followed by the correlation between dynamic and haptic (r^*2*^=.94, *p* <.01), and between static and haptic spaces (r^*2*^=.89, *p* <.01).

**Figure 4.**
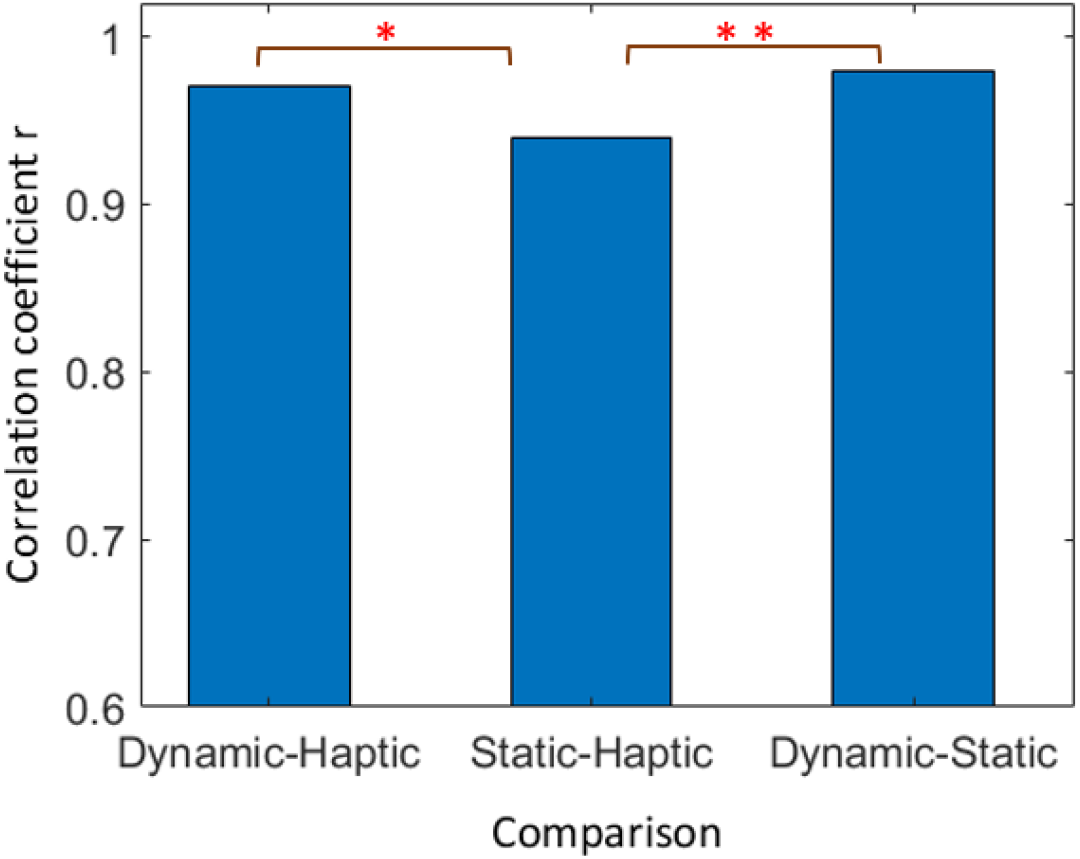
Comparisons of correlation coefficient *r* across dynamic visual information-haptic, static visual information-haptic, and dynamic visual information-static visual information conditions. Asterisks represent significance levels (*: *p* < 0.05, **: *p* <.01).

We next tested our two hypotheses, namely that the correlation between the two visual conditions should be significantly stronger than any other correlation and that the correlation between the dynamic visual condition and the haptic experiment should be stronger than the correlation between static visual condition and haptic experiment. The correlation between dynamic and static spaces was indeed significantly larger than that between static and haptic spaces (*p* < .01, one-tailed), however it was not significantly larger than the dynamic-haptic correlation. Pertaining to our second hypothesis we found indeed, that the correlation between the dynamic-haptic spaces was larger than that between static and haptic spaces (*p* =0.03, one-tailed).

It is further possible that the strength of the correspondence might vary between the respective softness dimensions, i.e., for *surface softness, granularity or viscosity*. To investigate this possibility, we computed the correlations of Bartlett scores also at the dimensional level. As expected, the correlations across conditions (static, dynamic, haptic) *within* the respective dimensions were very high and statistically significant (Figure 5, dark blue colors, all *p* <.01), and correlations across the respective dimensions were low and not significantly different from 0 (light blue colors).

**Figure 5.**
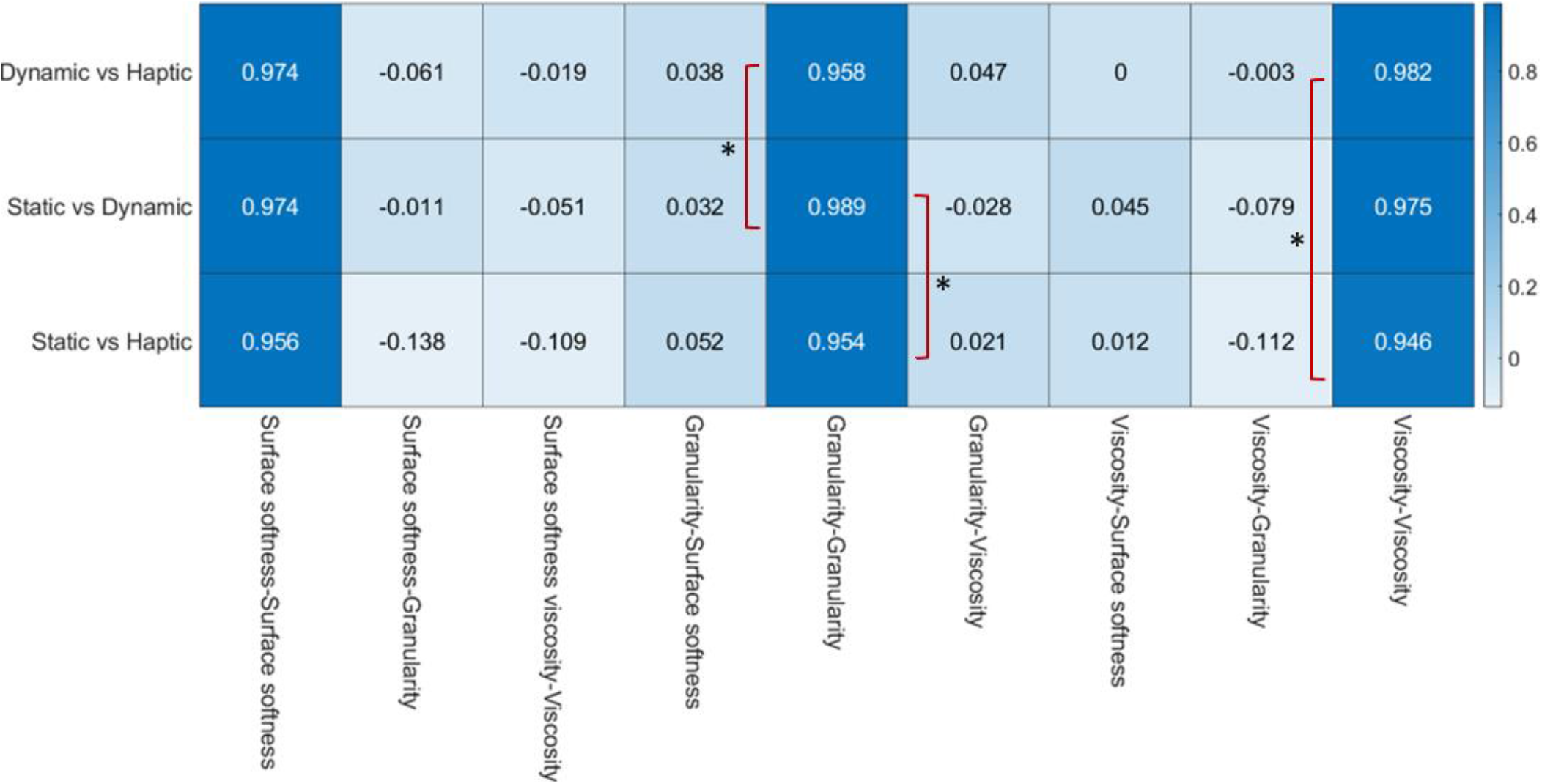
Correlations between Bartlett scores across materials for either pair of component scores from the haptic, the dynamic, and the static visual cue conditions. Darker colors (blue) show higher while lighter colors (white) show lower correlations.

We next put our two hypotheses to test. Regarding our first hypothesis, which states that the correlation between the two visual conditions should be strongest, we find that even though the correlation within *surface softness* between static-dynamic spaces was higher (.974) than that between static-haptic spaces (.956), this difference was not statistically significant. Within *granularity, however* the correlation between static-dynamic spaces (.989) was significantly higher (*p* =.01, one-tailed) than the correlation between static-haptic spaces (.954; *p* = .01, one-tailed), indicating a high correspondence between the two visual conditions for this dimension.Within the *viscosity* dimension the correlations between static-dynamic spaces (.975) was higher than the correlation between the static-haptic spaces (.946), however, this difference did not yield statistical significance.

As a reminder, our second hypothesis was that the correlation between the dynamic visual condition and the haptic experiment should be stronger than the correlation between static visual condition and haptic condition. While numerically this trend was true for all dimensions, only for *viscosity* the correlation between dynamic and haptic spaces (.982) was significantly higher than that between static and haptic spaces (.946, *p* = .03, one-tailed).

These analyses suggest that despite an overall good agreement between static, dynamic and haptic perceptual softness spaces, there are also some interesting differences as to how the softness of materials is represented in each of these spaces. In the next analysis we will analyze the rating differences of observers in the three conditions in order to understand for what material-adjective pairs the ratings of participants differ most.

### Rating differences between static, dynamic and haptic conditions

Overall, only 35 out of the 855 (285 × 3) rating differences exceeded the determined cut-off value. From these, three groups of difference patterns emerged: In one group there were *always* significant differences between haptic and the two respective visual conditions, but *never* between the two visual conditions. In Figure 6 we show these differences in a bar plot. The x-axis shows the corresponding adjective and material that elicited these rating difference. To appreciate what a specific bar height means, remember that ratings in all experiments varied between 1 (does not apply at all) and 5 (strongly applies). A positive difference implies that participants thought, that a particular adjective applied *more* to the material in question, e.g. that pebbles, lentils or cranberries felt more granular than they looked in either the dynamic or static conditions, that linen felt softer than it looked, or that fur feels more velvety than it looks. For this group, it appears that haptic information conveys information about material properties that is distinct from that conveyed by visual information, or conversely, the two visual conditions conveyed similar information (consistent with our 1^st^ hypothesis above).

**Figure 6.**
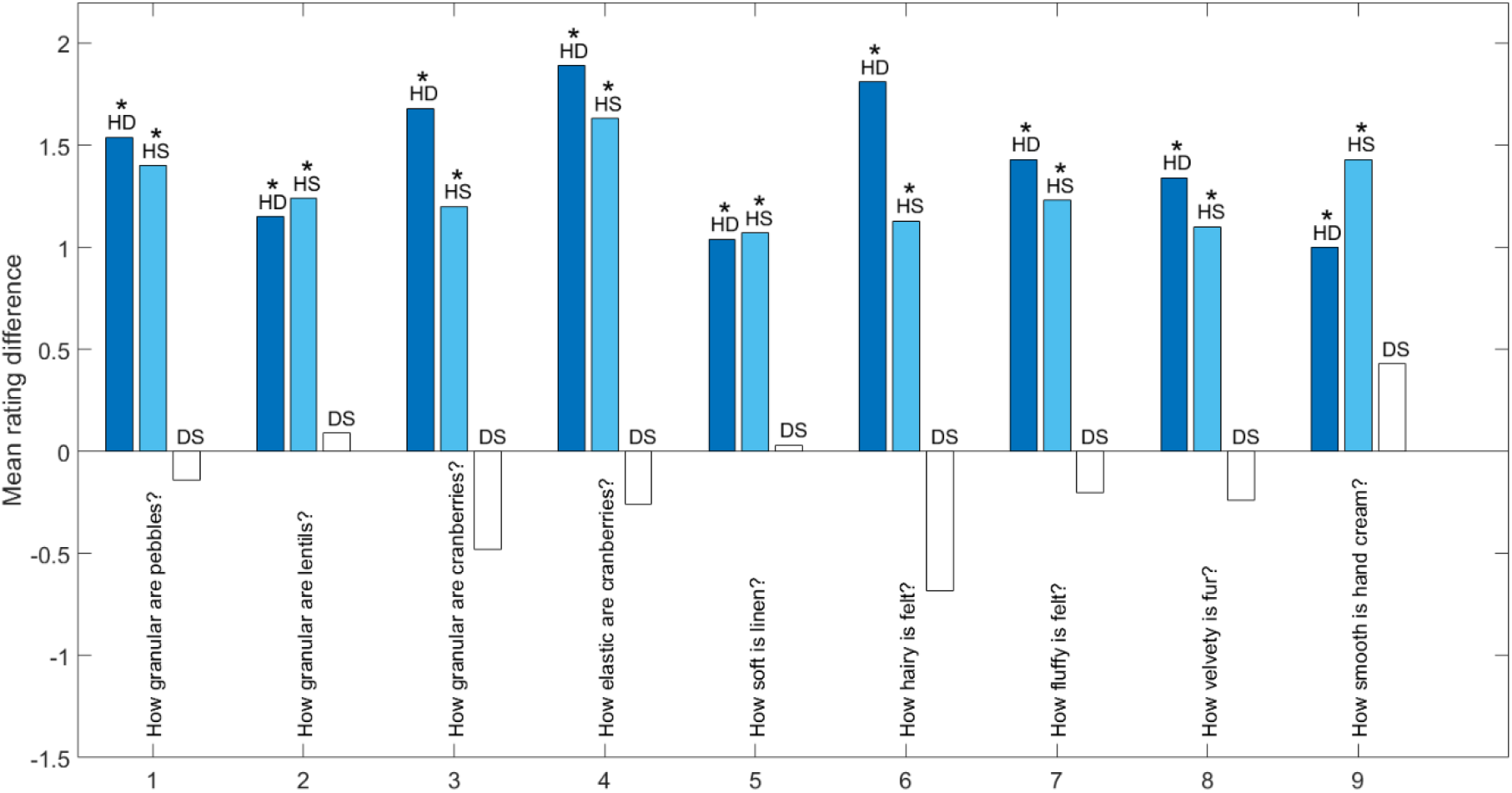
Haptic and visual information each convey different material qualities. Mean rating differences between static dynamic and visual conditions for specific material-adjective pairs. HD refers to the differences in rating between haptic and dynamic conditions (dark blue), HS to the differences in rating between haptic and static conditions (light blue), and DS to the differences in rating between dynamic and static conditions (white). * show the mean differences larger than 95% percentile cut off value (see Methods).

In a second group there was *never* a significant difference between haptic and dynamic conditions, but there was for other comparisons (e.g. for haptic-static, or for dynamic-static, Figure 7). For example, the softness of sand, salt and cranberries was judged equally in haptic and dynamic conditions, as was the hardness of stress balls. Here, it appears, that dynamic visual condition conveyed similar information as the haptic condition (as we also expected in our 2^nd^ hypothesis above). Additionally, the differences between the two visual conditions tended to be larger than in group one (Figure 6), even reaching significance in some cases.

**Figure 7.**
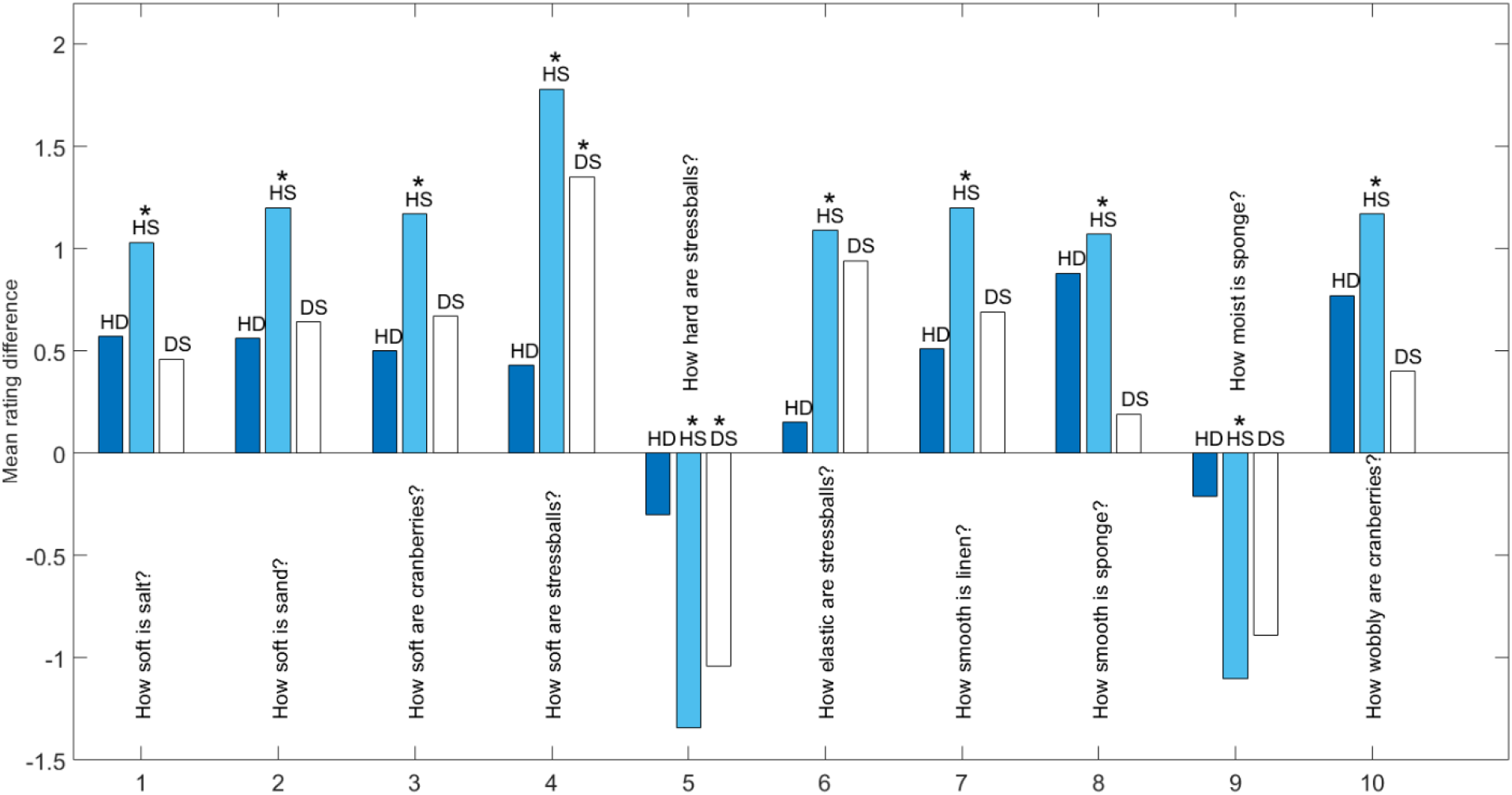
Dynamic visual and haptic information convey similar material properties. Mean rating differences between static dynamic and visual conditions for specific material-adjective pairs. Symbols and colors as in Figure 6.

Lastly, the third group of differences was the most surprising, containing cases with significant differences between haptic and dynamic conditions *only* (Figure 8). This goes directly against our 2^nd^ hypothesis, which proposed that dynamic and haptic conditions should yield more similar outcomes. Instead, for judgments of ‘how smooth lentils are’ or ‘how velvety linen is’, dynamic visual information appears to bias the participants away from the material properties perceived by inspecting a static image or by feeling the materials.

**Figure 8.**
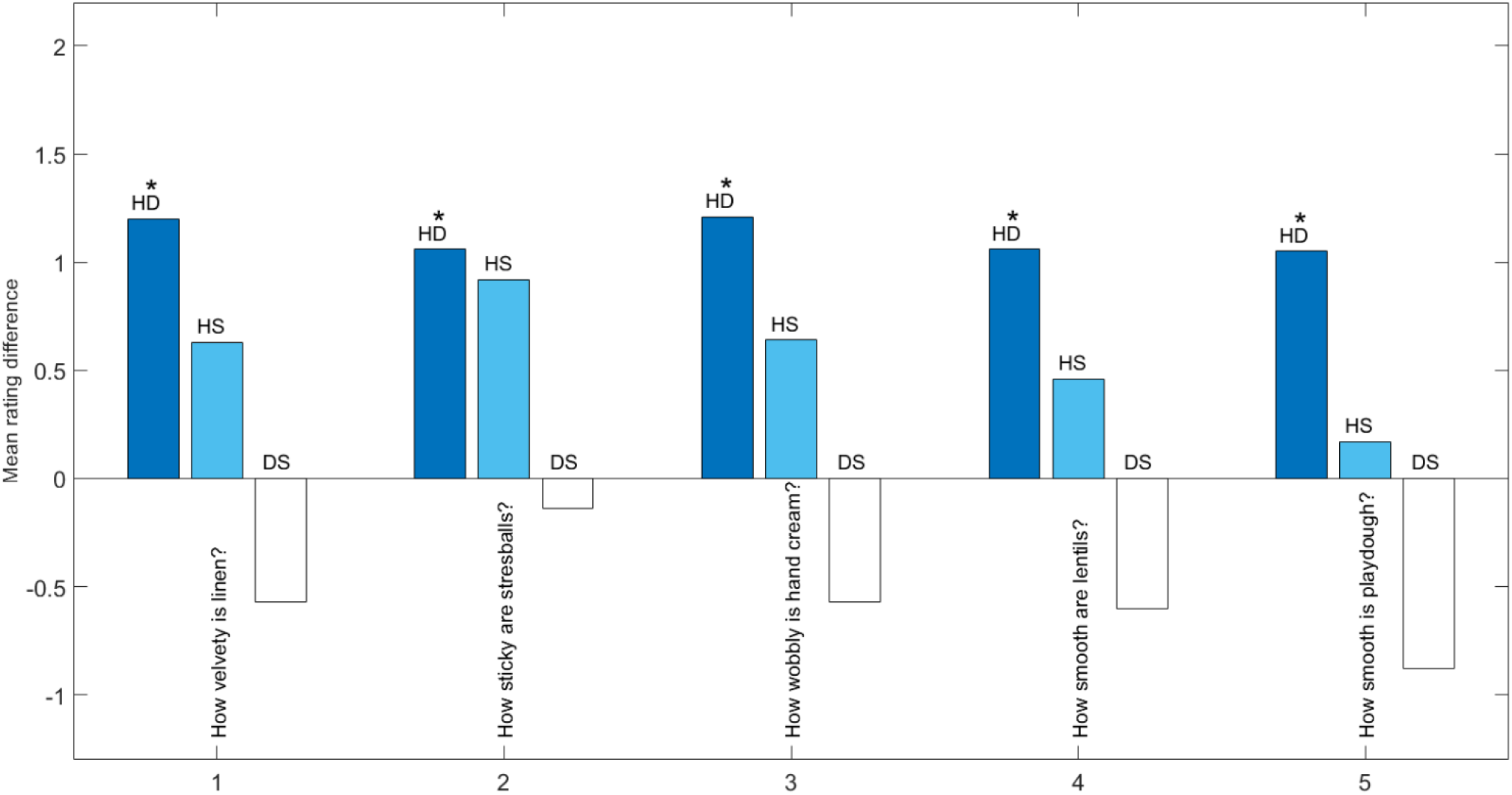
Static visual and haptic information convey similar material properties. Mean rating differences between static dynamic and visual conditions for specific material-adjective pairs. Symbols and colors as in Figure 6 and 7.

## Discussion

Softness is a prominent object property that renders it – depending on our intentions - useful (soft pillows) or useless (soft tables), appealing (soft fur) or repulsive (soft apples) to us. While we think of softness as primarily a mechanical property that can be perceived through touch (Klatzky & Lederman, 1987; Cellini et al., 2013; Okamoto et al., 2013; Di Luca, 2014; Higashi et al., 2019; Kitada et al., 2019; Cavdan et al., 2019; Dovencioglu et al., 2020; Xu et al., 2020) softness can also be judged visually (Drewing et al., 2009; Giesel & Zaidi, 2013; Baumgartner et al., 2013; Bouman et al., 2013; Bi & Xiao, 2016; Bi et al., 2018; Schmid & Doerschner, 2018). This latter ability is most likely acquired through countless multisensory interactions with objects in the environment, where simultaneous activation of visual and haptic senses leads to strong associations across modalities (Lacey et al., 2010; Yildirim & Jacobs, 2013; Desmarais et al., 2017). For example, while exploring a type of fabric (e.g. silk, or wool), its optical properties and the way it folds and deforms (i.e. its shape) might become associated with a particular perceived softness. This association can become so strong that when looking at an image of a material whose optical and shape properties strongly resemble the originally experienced fabric, it can elicit the same ‘sensation’ of soft (also see Schmidt et al. 2017; Anderson, 2011 or Schmid & Doerschner 2019, for a discussion of this potential association route). This might also explain why there is a high degree of consistence between visually tactile perceived material properties (Baumgartner et al., 2013; Vardar et al., 2019). However, to some degree this overlap is surprising, because of the inherently different information that is available in each sense. Whereas visual stimulus is basically a distal extended intensity pattern (image) that often changes across time (unless we look at a static image), haptic information is proximal, inherently serial, point by point and contains also direct signals about the applied force.

In this experiment we asked whether perceived softness from visual images and movies is comparable to perceived softness from haptic interactions (Cavdan et al., 2019). *The most important finding is that not just haptic, but also visually perceived softness is a multidimensional construct.* Consequently, one should keep this in mind when asking participants to judge the ‘softness’ of materials or objects in perceptual experiments. A second important result is that the haptic perceptual space is more differentiated (five dimensions) than the visual ones, with the dynamic visual space (four dimensions) resembling the haptic space more closely. Overall, we found beyond these differences in differentiation also very good agreement between the perceptual spaces yielded by visual and haptic experiments, which is also in line with earlier studies comparing texture perception across visual and haptic domains (Binns, 1936; Lederman & Abbott, 1981; Bergmann-Tiest & Kappers, 2007; Stilla & Sathian, 2008; Baumgartner et al., 2013; Vardar et al., 2019; Xiao et al., 2016).

In particular, we found three softness dimensions: *surface softness, granularity* and *viscosity* that were common to all conditions. While the amount of agreement between visual and haptic experiments is substantial for the softness dimensions of *surface softness, granularity* and *viscosity,* Table 1 also shows several interesting differences between these conditions, which we will review next.

### Differences in dimensionality

The individual principal component analyses revealed three softness dimensions: *surface softness, granularity* and *viscosity* in all three conditions (static, dynamic and haptic). However, in dynamic and haptic conditions also the dimension *deformability* emerged, and *roughness* emerged as a fifth dimension in the previous haptic experiment.

Why might *deformability* not have emerged as a separate dimension in the static condition? The deformability of a material is related to its kinematic properties and can therefore in static images only be judged from shape or texture cues (Schmidt et al., 2017; van Assen et al., 2018; Schmid & Doerschner 2017) or by association (Schmidt et al., 2017). Association, however, relies on two conditions: 1. the material has to be familiar and 2. the familiar material has had to be judged on the same attribute before. This might however not have been the case for many attribute material-combinations: participants might have never judged the elasticity of cranberries before and could thus not rely on their previous experience. Instead, they had to rely on the available image information (shape and texture cues) which might have highly overlapped with those used for surface softness. In contrast, dynamic visual information can convey the deformability of a material much more convincingly (Bouman et al., 2013; Bi & Xiao, 2016; Schmid & Doerschner, 2018, Schmidt et al., 2017, van Assen et al., 2018, Bi et al., 2018; Alley et al., 2020), in particular if also manual interactions with the material are shown (Cellini et al., 2013; Drewing & Kruse, 2014; Paulun et al., 2017; Yokosaka et al., 2018; Wijntjes et al., 2019).

Why might *roughness* not have emerged as a dimension in the visual conditions? In the present study there may have only been a limited number of adjectives that were strongly associated with the roughness dimension, namely smooth and rough. In haptics, roughness is a known as a particular salient dimension (e.g., Okamato et al., 2013), the value of which is quickly processed from the information gathered through the finger pads (Lederman & Klatzky, 1997). Thus, also with only limited measurement sensitivity, roughness can be detected as a haptic dimension. However, visually roughness is a much less salient and important dimension, and hence we might have missed to detect visually associated roughness in the present experiment. Indeed, visual ratings on roughness-related adjectives were not very variable across materials or used for dimensions other than roughness. Previous research on roughness perception found high correspondence between vision and touch (Brown, 1960; Björkman, 1967; for a review, see Lederman & Klatzky, 2004; Bergmann-Tiest & Kappers, 2007). However, tactile information, tended to be weighted more than visual information when the roughness information is mismatched between the two modalities (Guest & Spence, 2003; Whitaker et al., 2008; Eck et al., 2013), or while matching abrasive papers (Lederman & Abbott, 1981). Guest & Spence (2003) even reported a lack of visuo-tactile interactions for finer roughness stimuli. It could be that such fine texture information might have not been available in our visual conditions, which might explain the lack of a roughness dimension in the visual conditions. This would be consistent with the view that touch is superior to vision when detecting finer surface textures (Heller, 1989).

### Differences in the perceptual softness space structure

With a combined PCA we were able to zoom in on differences between static, dynamic and haptic spaces for the softness dimensions common to all three: *surface softness, granularity and viscosity*. As can be seen in Figure 4, the overall pattern that emerged when correlating the Bartlett scores between the three spaces (across all 3 dimensions) was that the two visual spaces were highly similar, however only when compared to the static-haptic correlation; dynamic-haptic spaces correlated just as high as the two visual spaces. However, we also noted some differences to this general pattern when looking at the Bartlett score correlation across spaces for each individual perceptual dimension, especially with respect to the latter finding. For example, while *surface softness* was numerically consistent with this general trend (Figure 5), there was no significant difference in the correspondence between spaces. For *viscosity*, on the other hand, the correlation between dynamic and haptic spaces was significantly stronger than that between dynamic and static spaces. What might be the reason for this? Inspecting, Figure 3, might provide a hint: the Bartlett scores of the material *stress balls* show high values in *viscosity* of haptic and dynamic spaces but not in the static space. *Stress balls,* although being quite squishy and sticky to the touch, do in their ‘resting’ shape not convey these properties strongly. Therefore, the shape of the material might cause the visual system to activate a not-so-viscous material association (Schmidt et al., 2017). This emphasizes the high relevance of dynamic visual information in transporting viscosity. While it is undebated that static images *can* successfully convey information about viscosity, they do so primarily via (shape) association. However, when the shape is unfamiliar or unusual static images will not be able to unambiguously convey the viscosity of a material.

Another exception to the described overall correlation pattern was found for *granularity*. Here, the correspondence between the two visual conditions was significantly higher than between dynamic and haptic spaces. This suggests two things: 1. Granularity can be judged well and consistently from images, with observers likely using the size of individual items (sand corns, lentils, pebbles etc.), which would be available both in static images and videos. 2. These visually estimated properties differ from those estimated by touch. This might be because vision strongly relies on particle size while touch might additionally consider interaction characteristics of the particles (e.g. how well they can be run through the fingers or be rotated).

### Differences in ratings

Our interpretations above suggest that the differences that we find between the perceptual softness spaces of static, dynamic and haptic conditions might be particularly driven by some special material-adjective combinations in our experiments. In order to sift these out we identified the conditions that yielded the largest rating differences across conditions. Only 35 rating differences survived this procedure, again supporting the idea that perceived softness appears to be similar between visual and haptic spaces. Those material-adjective combinations that yielded significant rating differences between conditions fell into three categories: 1) haptic and visual information each convey different material qualities 2) dynamic visual and haptic information convey similar material properties, 3) static visual and haptic information convey similar material properties. We will discuss each one of these next.

#### Haptic and visual information each convey different material qualities

For the material adjective combinations in this group, it appears that haptic information conveys information about material properties that is distinct from that conveyed by visual information, or conversely, the two visual conditions conveyed similar information. Looking at the specific cases for which this occurs we see that this pattern emerged primarily for judgments related to the granularity (how granular and surface softness (how hairy, velvety, hairy). We already offered an explanation about the differences between visual and haptic perception of granularity above (vision relies on particle size when touch might rely on interaction characteristics of the particles, see section ‘differences in the perceptual softness space structure’). Why do we, however not see such differences for granularity judgments of sand. It is possible sand or salt are materials that most observers are very familiar with, and when identifying the materials, memories of interacting with the material might become activated enabling participants to make these judgments. For example, found that perceived softness in haptic experiments is influenced by memory (Metzger & Drewing, 2019), and haptic experiences (Kangur et al., 2019). Conversely, it is possible that, when judging the *granularity* of lentils, pebbles, or cranberries, such a prior experience is not available and therefore, participants are left with visual information ‘only’, which might lead to different perceptions.

This kind of argument could also be made for judgments of *surface softness*. Figure 6 shows that the differences between visual and haptic conditions are generally positive suggesting that felt fur and linen were judged *more* soft, hairy, and velvety, respectively when interacted with. In a sense, the experiences of *surface softness* tend to be lower from visual images, which highlights the special role of interactive touch for perceiving this material quality.

#### Dynamic visual and haptic information convey similar material properties

In contrast to the first group, adjective-material pairs in this group elicited similar judgments in dynamic and haptic experiments, whereas the difference to ratings in the static condition increased (height of white bars in Figure 7). Why did static visual information yield different ratings than haptic or dynamic conditions? Figure 7 illustrates that this kind of pattern emerged primarily for judgments of *surface softness* (how soft), but also for judgments of *deformability* (how elastic, how hard), or *viscosity* (how moist, how wobbly). The fact that surface softness occurs also in this group is unexpected, as we have just concluded that softness tends to be lower from any type of visual information, compared to that from haptic experience. It appears that we will have to modify this statement. How can we reconcile the data from Figures 6 and 7? Let’s consider the stimuli in the dynamic condition: the movies contained three sets of cues to material properties 1) pictorial cues, 2) deformation cues and 3) interaction cues. While pictorial cues must have played a predominant role in the difference pattern of the first group (i.e., haptic and visual information each convey different material qualities), we believe it is the second set of cues that might be responsible for the higher similarity between dynamic and haptic ratings. We gave already a concrete example along this line with the elasticity (or hardness) rating of stress balls in the section above, but similar arguments can be made for judging the wobbliness of cranberries or the softness of sand. Why deformation were not effective for the first group of material-adjective pairs, would be an interesting question to pursue in future research.

#### Static visual and haptic information convey similar material properties

Although this occurred just for 5 material-adjective pairs, this was the most interesting pattern, as it was in contradiction to our hypothesis, that dynamic and haptic conditions should yield more similar outcomes (i.e. the pattern observed in group 2). Why might haptic and static conditions yield more similar ratings? One interpretation is that static images triggered associations of material qualities that were similar to those experienced through haptic exploration. It is possible that – in contrast - dynamic stimuli, showed either deformation of interaction cues that elicited a slightly different activation of material properties. Why might this be the case? We have shown, for example, in previous work (Cavdan et al., 2019, 2020) that exploratory hand movements not only vary with the material being explored but also as a function of the task (i.e., what is being judged while exploring a material). When selecting the stimuli for the dynamic condition we focused on the most frequent hand movement for a material type, neglecting the effect of task (since it was a smaller affect in our previous work). For example, for lentils we only used the hand movement *run through*, yet people might need to see *rotation* in order to understand how smooth lentils are. It might be that this very subtle factor might have influenced observers’ judgments in the dynamic condition. This possibility could be explored in future work.

## Conclusion

Softness is a prominent property that renders an object useful or useless, appealing or repulsive to us. This study shows that perceived softness is a multidimensional construct, and this should be taken into consideration when asking participants to make judgments about the softness of materials in research or applied contexts. This multidimensional softness space is similar for visually and haptically presented materials, however, we also found some noteworthy differences. We argue that these differences appear primarily to emerge when participants cannot draw on previous visuo-haptic experiences with a material for a particular judgment, or when visual cues are ambiguous to the material property in question.

## Footnotes

1 Also, auditory cues play a role in material perception, however these are not the focus of this investigation (Klatzky et al., 2000; Fujisaki et al., 2014; Fujisaki et al., 2015).

## Acknowledgement

This research was supported by the EU Marie Curie Initial Training Network “DyVito” (H2020-ITN, Grant Agreement: 765121). Further support was given by a Sofja Kovalevskaja Award by the Alexander von Humboldt Foundation endowed by the by Deutsche Forschungsgemeinschaft (DFG, German Research Foundation), and an SFB/TRR 135, A5. The authors thank Maj Charlotte Brinkmann and Anna-Klara Trieschmann for data collection and Robert Ennis for technical support.

