## Supplemental File 1 for "Materials in action: The look and feel of soft"

### Supplementary Materials

#### Supplementary Methods 1

Summary of selected exploratory movements from Dovencioglu et al., 2020 and Cavdan et al., 2019:

1. Run through: "Picking up some parts/portion of the material and letting them trickle through the fingers."

2. Rotate: "Lifting parts of the material to move and turn its boundaries typically inside the finger(tip)s."

3. Rub: "Applying torque or lateral force with varied pressure levels, sometimes sweeping materials between index and thumb fingers or forcefully stroking material with the thumb while poising the object with the other fingers."

4. Pull: "Stretching a part of the material by moving fingers or separating them from each other."

5. Pressure: "Applying directional normal force to squeeze a material between palm and fingers or using one or more fingers to apply normal force."

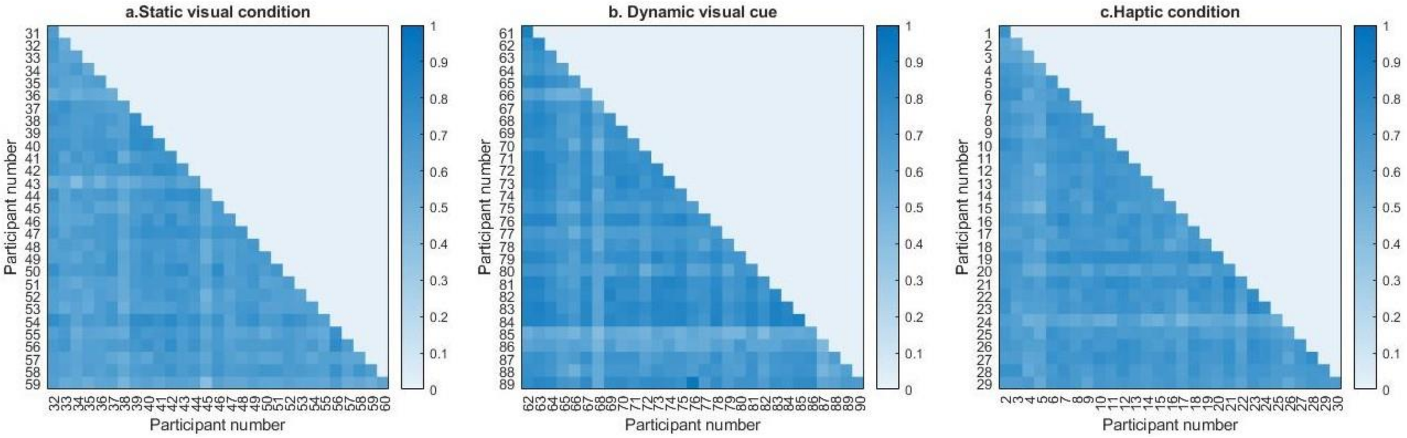


**Supplementary Figure 1**. **Consistency across participants.** Correlation coefficients across participants for (a) static (b) dynamic (c) haptic conditions (Pearson's correlation for each pair of participants across 19 materials times 15 adjective ratings). High correlations are plotted in dark blue and low correlations are plotted in light blue (white would be $r = 0$). Correlations of Ratings across participants were ranging between .45 to .86, .41 to .81, and .41 to .95, sequentially for haptic, static visual information, and dynamic visual information conditions and all were statistically significant at $p < .01$.

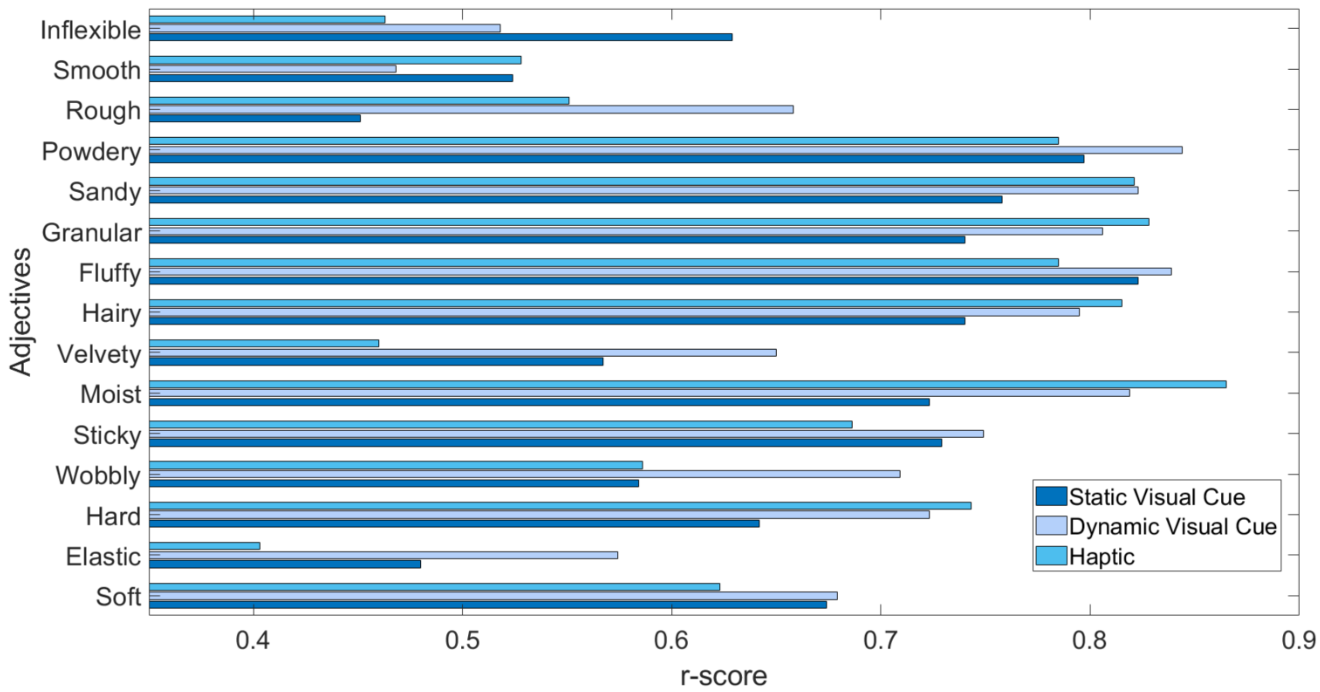


**Supplementary Figure 2. Mean inter-observer consistency per adjective**. Mean correlation coefficient across materials and participants per adjective in static, dynamic, and haptic conditions. Mean correlations range from.45 to .82, .47 to.84, and .40 to .86, in static, dynamic and haptic conditions, respectively.

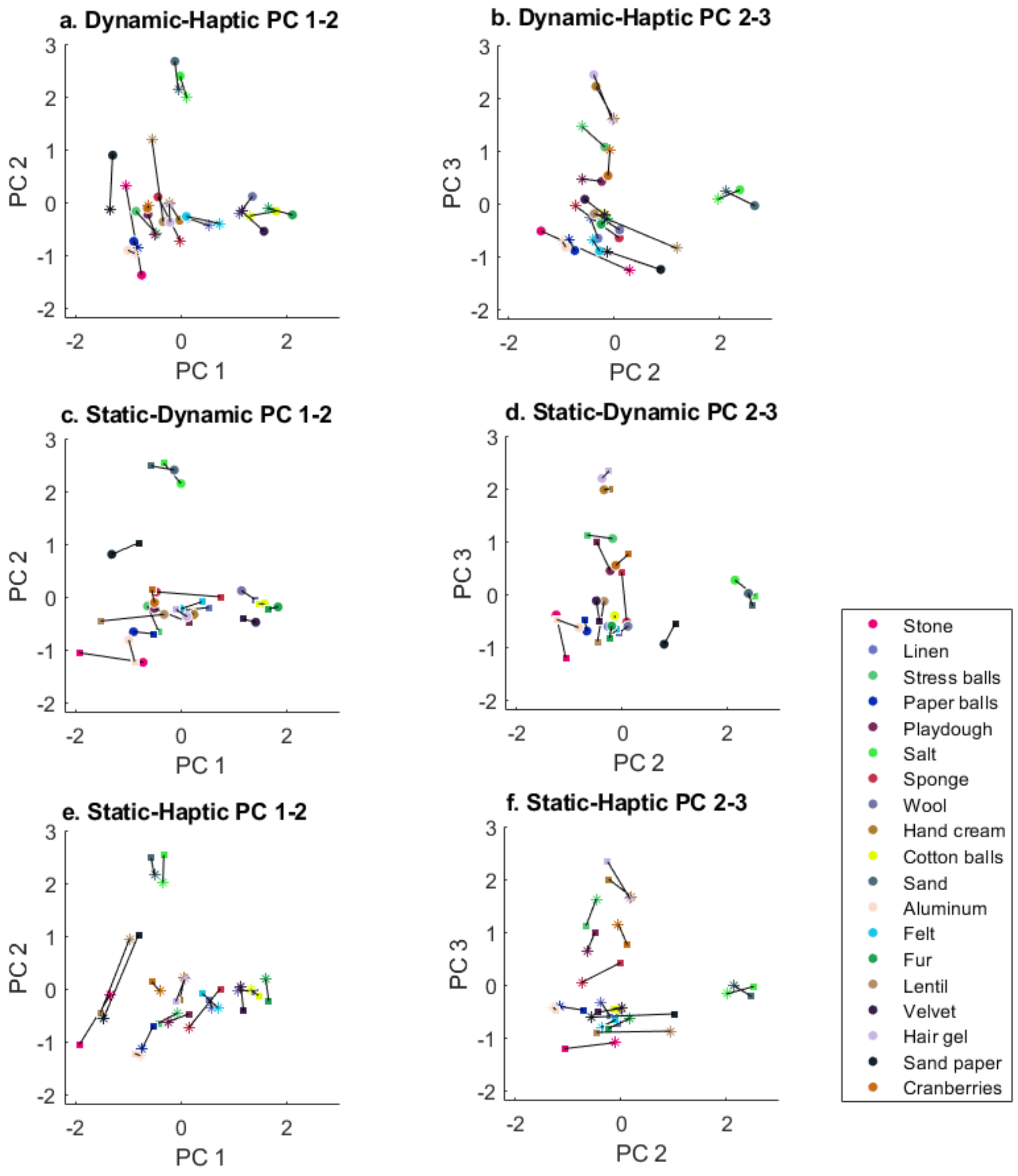


**Supplementary Figure 3.** Procrustes analysis of the change in Bartlett score values of each material across the three perceptual softness dimensions: surface softness, granularity, viscosity when mapping from dynamic to haptic (a. dynamic-haptic principal component 1 vs 2, b. dynamic-haptic principal component 2 vs 3), static to dynamic (c. static-dynamic principal component 1 vs 2, d. static-dynamic principal component 2 vs 3 ), and static to haptic (e. static-haptic principal component 1 vs 2, f. static-haptic principal component 2 vs 3). Each symbol represents one condition (dot: dynamic, square: static, and star: haptic). The second title condition indicates the transformed spaces.
